# The seminal vesicle is a juvenile hormone-responsive tissue in adult male *Drosophila melanogaster*

**DOI:** 10.1101/2024.03.20.585833

**Authors:** Yoshitomo Kurogi, Yosuke Mizuno, Naoki Okamoto, Lacy Barton, Ryusuke Niwa

**Author notes:** Correspondence: Ryusuke Niwa.

## Abstract

Juvenile hormone (JH) is one of the most essential hormones controlling insect metamorphosis and physiology. While it is well known that JH affects many tissues throughout the insects life cycle, the difference in JH responsiveness and the repertoire of JH-inducible genes among different tissues has not been fully investigated. In this study, we monitored JH responsiveness *in vivo* using transgenic *Drosophila melanogaster* flies carrying a *JH response element-GFP* (*JHRE-GFP*) construct. Our data highlight the high responsiveness of the epithelial cells within the seminal vesicle, a component of the male reproductive tract, to JH. Specifically, we observe an elevation in the JHRE-GFP signal within the seminal vesicle epithelium upon JH analog administration, while suppression occurs upon knockdown of genes encoding the intracellular JH receptors, *Methoprene-tolerant* and *germ cell-expressed*. Starting from published transcriptomic and proteomics datasets, we next identified *Lactate dehydrogenase* as a JH-response gene expressed in the seminal vesicle epithelium, suggesting insect seminal vesicles undergo metabolic regulation by JH. Together, this study sheds new light on biology of the insect reproductive regulatory system.

## Introduction

Juvenile hormone (JH) was initially discovered in the 1930s as an insect metamorphosis inhibition factor [1–4]. JH is synthesized in the *corpora allata* (CA) and regulates many aspects of insect physiology throughout the life cycle [5–8]. JH signaling is mediated through intracellular JH receptors, Methoprene-tolerant (Met) and its paralogs, which belong to the basic helix-loop-helix (bHLH)-Per-Arnt-Sim (PAS) family of transcriptional factors [9–12]. Met and its paralogous transcription factors bind to JH with high affinity [10,13]. Upon JH binding, these intracellular receptors associate with specific JH response elements (JHREs), containing a C-box sequence (CACGCG, an E-box-like motif) or a canonical E-box sequence (CACGTG) [13], followed by the transcriptional induction of target genes, such *Krüppel-homolog 1* (*Kr-h1*) [13–18].

In the last decade, the fruit fly *D. melanogaster* has contributed to elucidating molecular mechanisms of JH-responsiveness [19]. Two intracellular JH receptors have been identified in *D. melanogaster*, known as Met and Germ cell-expressed (Gce). Single loss-of-function of either *Met* and *gce* is adult viable, while double mutants of *Met* and *gce* result in developmental arrest during pupation, like CA-ablated flies [9,20], suggesting that Met and Gce act redundantly to regulate JH-responsive gene expression. A recent study using *GAL4*– and *LexA*-based reporters showed that *Met* and *gce* are both broadly expressed in many, but not all, tissues throughout *D. melanogaster* development [21], suggesting many tissues have the potential to transcriptionally respond to JH. Yet, whether all tissues that express JH receptors have active JH transcriptional signaling is unknown.

To approach this problem, we conducted a study using a *D. melanogaster* strain carrying a *JH response element-GFP* (*JHRE-GFP*) [22]. The *JHRE-GFP* construct contains eight tandem copies of a JHRE, originally identified from the *early trypsin* gene of *Aedes aegypti* [16,17,23]. It has also been confirmed that *JHRE* is responsive to JH analogs in *D. melanogaster* S2 cultured cells [10]. In addition, a recent study has shown that GFP signals of *JHRE-GFP* transgenic flies can monitor JH-responsiveness in *D. melanogaster* embryos [22]. In this study, we show that JHRE-GFP signal is found in epithelial cells of the adult seminal vesicle, which is a part of the male reproductive tract in *D. melanogaster*. The JHRE-GPF signal in the seminal vesicle epithelium is elevated upon JH analog administration and conversely suppressed in animals depleted of *Met* and *gce* by RNAi. We also show that JHRE-GFP in the seminal vesicle epithelium is elevated after mating, consistent with a previous hypothesis that mating elevates JH titer in male adults [24,25]. Furthermore, we identified *Lactate dehydrogenase* (*Ldh*) as a JH-response gene expressed in the seminal vesicle epithelium. Our study demonstrates the seminal vesicle as a novel JH-responsive tissue in *D. melanogaster*.

## Results

### The seminal vesicle in male *D. melanogaster* is a JH-responsive tissue

In previous studies, while the functions of JH during development and its effects on the reproductive system of adult females have been extensively studied [1–8,19], its functions in adult males have received less investigation. Therefore, we investigated which cells/tissues are responsive to JH in the adult males using *JHRE-GFP* transgenic flies. Whereas *JHRE-GFP* strain has been used for monitoring JH-responsive cells during embryogenesis [22], it has not been used for adult males. Therefore, we first examined JHRE-GFP fluorescence signals in whole male adult bodies. We used two strains in this study, namely *JHRE^Wild-type^ ^(WT)^-GFP* males with *JHRE^Mutated^ ^(Mut)^-GFP* males [22]. *JHRE^WT^-GFP* strain carries a wild-type JHRE, while *JHRE^Mut^-GFP* strain carries a mutated JHRE in which Met and Gce binding sites are disrupted [10,22]. We observed strong GFP signals in the scattered hemocytes and some tissues in the abdominal region of *JHRE^WT^-GFP*, but not *JHRE^Mut^-GFP* flies (figure 1*a*). We also orally administrated a JH analog (JHA), methoprene, to these animals, and found that the GFP signals in the abdomen were particularly elevated in *JHRE^WT^-GFP*, but not *JHRE^Mut^-GFP* flies (figure 1*a*, arrowhead). Based on the data, we further anatomically characterized where *JHRE^WT^-GFP* was expressed in abdominal tissues.

**Figure 1.**
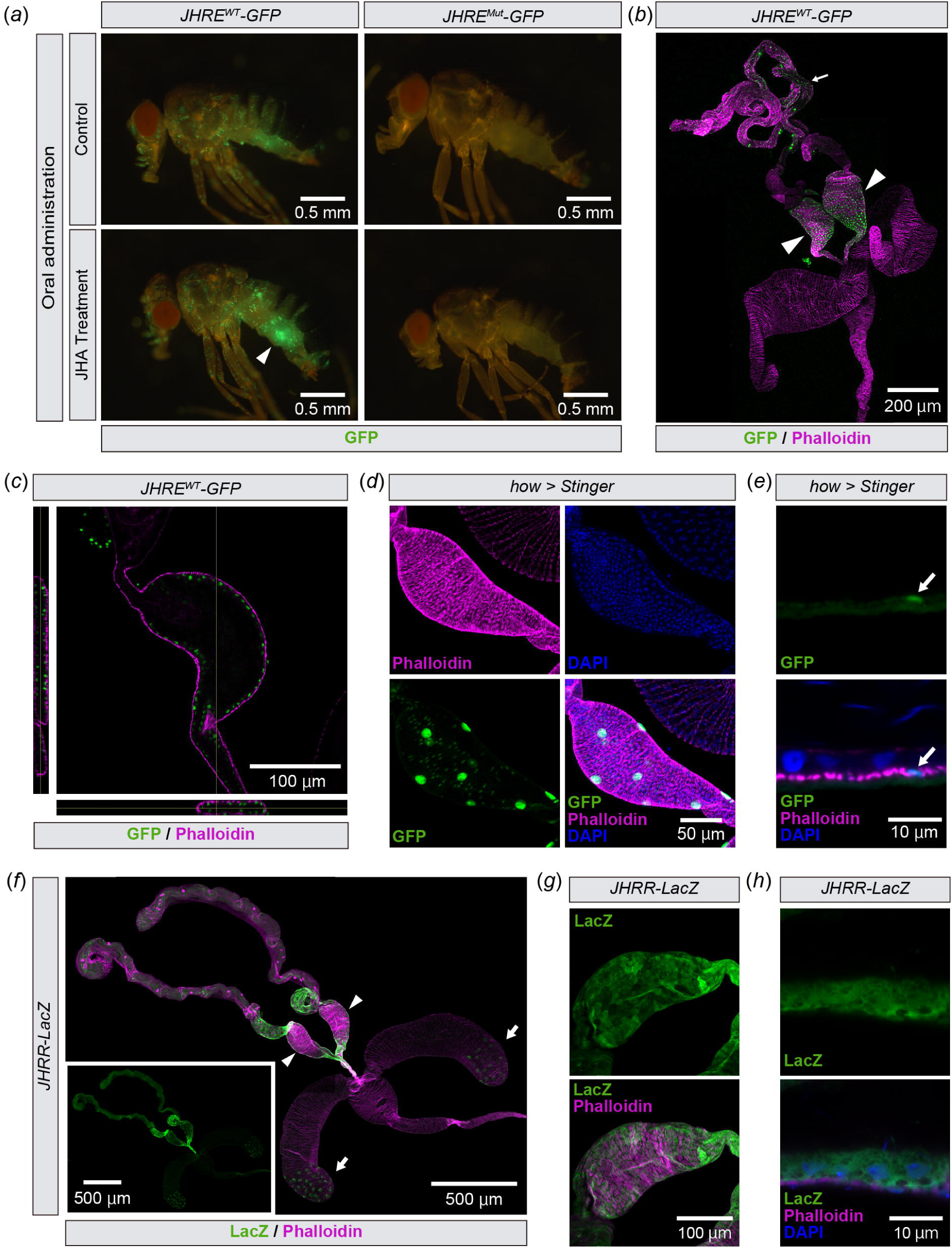
*JHRE-GFP* is expressed in seminal vesicle epithelial cells. (*a*) Whole body image of *JHRE^WT^-GFP* (left) and *JHRE^Mut^-GFP* virgin males (right) 7 days after eclosion, with or without oral administration of methoprene (JHA). GFP signals (green) in the abdomen of *JHRE^WT^-GFP* were increased by JHA administration (arrowhead). (*b*, *c*) Immunostaining with anti-GFP (green) and phalloidin (magenta) of *JHRE^WT^-GFP* adult male. (*b*) Image of the male reproductive tract. The arrowhead indicates the seminal vesicles. (*c*) Cross-section image of the seminal vesicle. Left and bottom images indicate horizontal and vertical cross-sectional views, respectively. (*d*, e) Transgenic visualization of muscles by nuclear GFP (Stinger) driven by *how-GAL4*. Samples were immunostained with anti-GFP antibody (green), phalloidin (magenta) and DAPI (blue). Samples were derived from virgin males 2 days after eclosion. (*d*) Image of the seminal vesicle. (*e*) Magnified view of the seminal vesicle epithelial cells. (*f*-*h*) Immunostaining with anti-LacZ (green) and phalloidin (magenta) of *JHRR-lacZ* adult virgin males 4 days after eclosion. (*f*) Image of the male reproductive tract. Arrowheads and arrows indicate the seminal vesicles and the male accessory glands, respectively. (*g*) Image of the seminal vesicle. (*h*) Magnified view of the seminal vesicle epithelial cells. Blue is the DAPI signal.

Dissection of *JHRE^WT^-GFP* male abdomens revealed that the JHRE-GFP signal was present in a part of the male reproductive tract, including the testes and seminal vesicles (Figure 1*b*), which is known to store sperm produced in the testis [26]. As the seminal vesicle showed the most remarkable JHRE-GFP signal in the male reproductive tract, we decided to focus on this tissue for the rest of this study. Within the seminal vesicles, JHRE-GFP was active in cells located on the lumen side compared to the muscle layer surrounding the seminal vesicle labeled with fluorescence-conjugated phalloidin (figure 1*c*). We assume that these luminal side cells were not muscle cells but epithelial cells, as *GFP* driven by the muscle driver *how-GAL4* [27] was expressed in fewer cells than JHRE-GFP-positive cells in the seminal vesicles (figure 1*d*) and embedded in the phalloidin-positive muscle layer (figure 1*e*). These results suggest that *JHRE-GFP* is expressed in the seminal vesicle epithelial cells.

We also examined whether these cells were labeled with another JH reporter strain, *JH response region* (*JHRR*)*-lacZ*. *JHRR-lacZ* is a *lacZ* reporter fused with the JHRR of the *D. melanogaster Kr-h1* promoter, which is responsive to JH via Met and Gce [14]. We found that *JHRR-lacZ* was also expressed in the seminal vesicle cells, some cells in the testes just anterior to the seminal vesicle, and some secondary cells of the male accessory gland (figure 1*f*). Similar to *JHRE-GFP*, *JHRR-lacZ* also labeled the epithelial cells of the seminal vesicles (figure 1*g*, *h*). These results support the idea that seminal vesicle epithelial cells are sensitive to JH.

We next examined whether seminal vesicle cells respond to the JH signaling. We found that the oral administration of JHA increased the JHRE-GFP signal in the seminal vesicles of virgin males carrying the *JHRE^WT^-GFP*, but not *JHRE^Mut^-GFP*, transgene (figure 2*a*,*b*). In addition, JHRE-GFP signal was elevated in *ex vivo* cultured seminal vesicles 16 hr after incubation with JHA (figure. 2*c*,*d*), suggesting that the seminal vesicle itself responds to JH. Conversely, when JH biosynthesis was blocked by knocking down *juvenile hormone acid O-methyltransferase* (*jhamt*), a rate-limiting enzyme for JH biosynthesis in the CA [28,29], JHRE-GFP signal in the seminal vesicle decreased (figure 2*e*,*f*). These results suggest that the seminal vesicle epithelial cells respond to changes in circulating JH.

**Figure 2.**
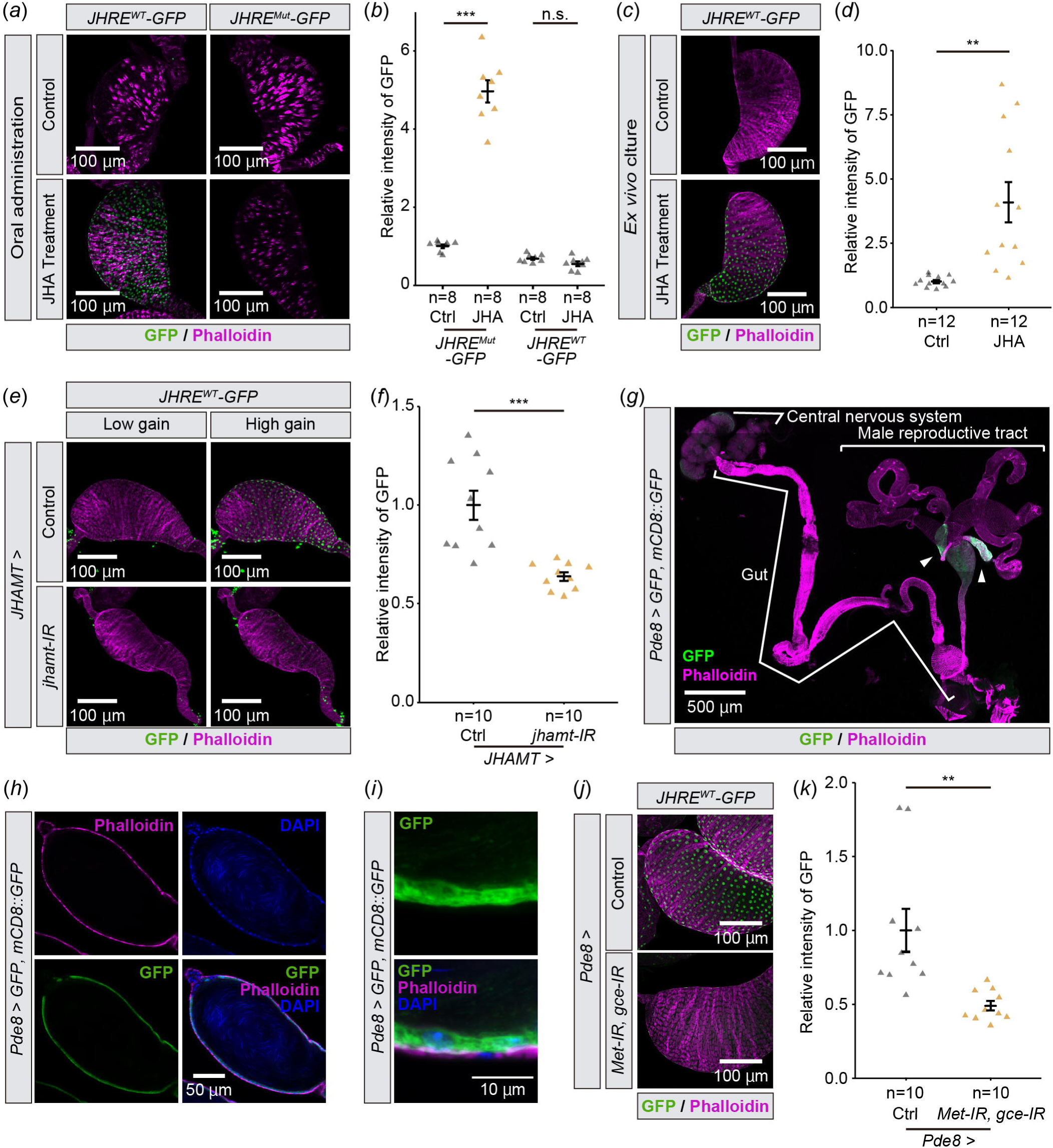
JHRE-GFP signal in the seminal vesicle changes depending on JH signaling. All samples were obtained from virgin males. In all photos, GFP and phalloidin (F-actin) signals are shown in green and magenta, respectively. (*a*, *b*) JHRE-GFP signal in the seminal vesicle of *JHRE^WT^-GFP* (left) and *JHRE^Mut^-GFP* males (right) 7 days after eclosion, with or without oral administration of methoprene (JHA). (*a*) Representative images of the seminal vesicles. (*b*) Quantification of JHRE-GFP signals in the seminal vesicles of control (Ctrl) and JHA-administrated (JHA) males. (*c*, *d*) JHRE-GFP signals in the seminal vesicle of *JHRE^WT^-GFP* males 4 days after eclosion. Male reproductive tracts without male accessory glands are *ex vivo* cultured with (Ctrl) or without methoprene (JHA). (*c*) Representative images of the seminal vesicles. (*d*) Quantification of JHRE-GFP signal in the seminal vesicles. (*e*, *f*) JHRE-GFP signal in the seminal vesicle of control and *JHAMT-GAL4*-driven *jhamt* RNAi males 4 days after eclosion. Control RNAi was achieved with a VDRC KK control line. (*e*) Representative images of the seminal vesicles. “Low gain” GPF signals were captured with the same gain as shown in (*c*). “High gain” GPF signals were captured with 1.23-fold gain setting compared with “Low gain” (800 vs 650). (*f*) Quantification of JHRE-GFP signal in the seminal vesicle. (*g*-*i*) Immunostaining with anti-GFP, phalloidin, and DAPI (Blue) of *Pde8-GAL4 UAS-GFP UAS-mCD8::GFP* males 4 days after eclosion. (*g*) Image of the central nervous system, gut, and male reproductive tract. Allow heads indicate the seminal vesicles. (*h*) Cross section image of the seminal vesicle. (*i*) Magnified view of the seminal vesicle epithelial cells. (*j*, *k*) JHRE-GFP signal in the seminal vesicle of control males and *Pde8-GAL4*-driven *Met* and *gce* RNAi males 7 days after eclosion. Note that this experiment was conducted with food supplemented with JHA, as the JHA administration allowed us to see more drastic difference in JHRE-GFP signals between control and RNAi. (*j*) Representative images of the seminal vesicles. (*k*) Quantification of JHRE-GFP signal in the seminal vesicle. Values in *b*,*d*,*f* and *k* are presented as mean ± SE. Statistical analysis: Student’s t-test for *b*,*d*,*f* and *k*. \*\**P* <0.01 \*\*\**P* <0.001. n.s.: not significant.

### JH signaling in the seminal vesicle requires Met and Gce

We next confirmed that *JHRE-GFP* expression in the seminal vesicle was mediated by intracellular JH receptors, Met and Gce [13–18]. However, since *Met* and *gce* double mutant flies die during the larval-pupal transition [9], we conducted transgenic RNAi to knockdown *Met* and *gce* with a GAL4 driver that labels the seminal vesicle epithelial cells. After our GAL4 driver screen (See Materials and Methods for details), we found that *Pde8-GAL4* driver drives gene expression in the seminal vesicles (figure 2*g*). Our further detailed analysis confirmed that *Pde8-GAL4* labels the seminal vesicle epithelial cells (figure 2*h*,*i*). Using this GAL4 driver, we found that JHRE-GFP signal in the seminal vesicle epithelial cells was decreased by *Met* and *gce* double knockdown (figure 2*j*,*k*). These results suggest that JH is received by Met and/or Gce in the seminal vesicle epithelial cells.

### Mating activates JH signaling in the seminal vesicle

Next, we tested whether *JHRE-GFP* expression in the seminal vesicle is responsive to natural processes reported to impact JH signaling. In *D. melanogaster* males, JH signaling may increase in a mating-dependent manner [24,25]. These previous observations motivated us to compare JHRE-GFP signals in the seminal vesicle between virgin and mated males. We found that JHRE-GFP signal in the seminal vesicle epithelial cells increased in mated males as compared to virgin males (figure 3*a*,*b*). In addition, the increase of the JHRE-GFP signal upon mating was canceled by *jhamt* RNAi in the CA (figure 3*c*,*d*). These results suggest that JH signaling in the seminal vesicle epithelium is responsive to mating.

**Figure 3.**
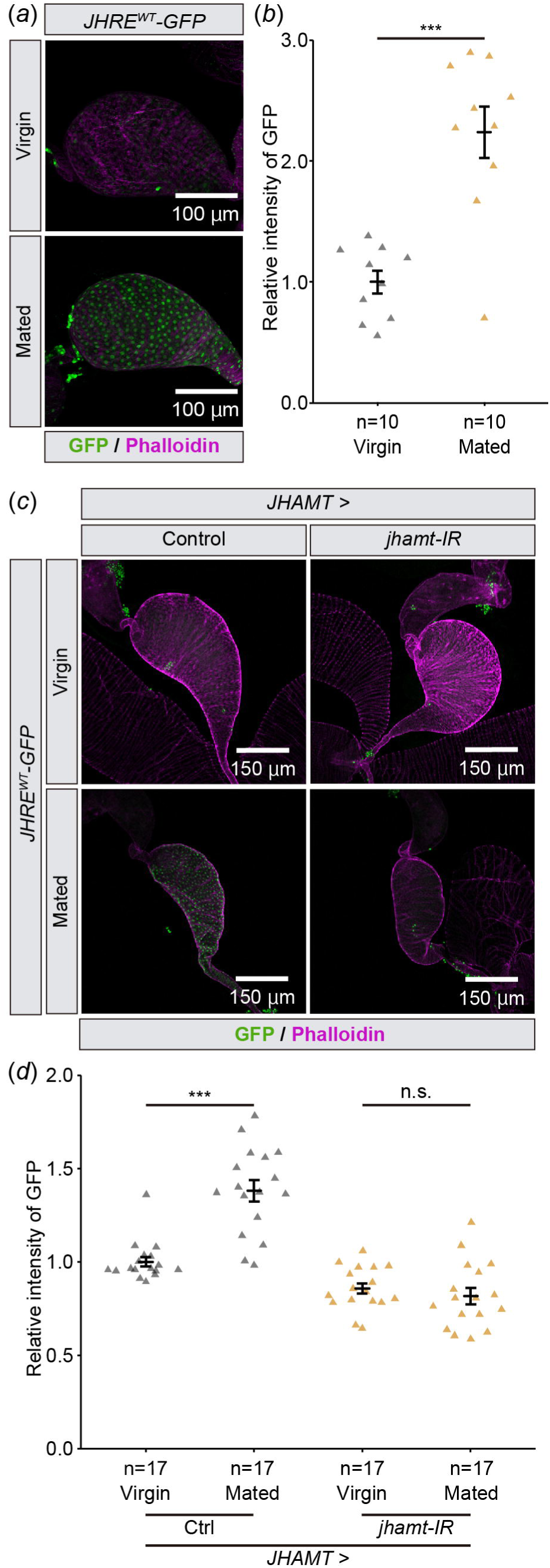
JHRE-GFP signal in the seminal vesicle is increased after mating. Samples were derived from males 6 days after eclosion. In all photos, GFP and phalloidin (F-actin) signals are shown in green and magenta, respectively. (*a*, *b*) JHRE-GFP signals in the seminal vesicle of virgin or mated males. (*a*) Representative images of the seminal vesicles. (*b*) Quantification of JHRE-GFP signals in the seminal vesicles. (*c*, *d*) JHRE-GFP signals in the seminal vesicles of control and *JHAMT-GAL4*-driven *jhamt* RNAi males with or without mating. Control RNAi was achieved with VDRC KK control line noted in the Methods section. (*c*) Representative images of the seminal vesicles. (*d*) Quantification of JHRE-GFP signals in the seminal vesicles. Values in *b* and *d* are presented as mean ± SE. Statistical analysis: Student’s t-test for *b*. Tukey-Kramer test for *d*. \**P* <0.05, \*\*\**P* <0.001. n.s.: not significant.

### JH induces expression of *Lactate dehydrogenase* in the seminal vesicle

In JH-responsive cells/tissues, JH signaling affects gene expression via Met and Gce [6,13]. Therefore, we searched for genes that are highly expressed in the seminal vesicles and potentially regulated by JH. First, we listed genes that might be highly expressed in the seminal vesicles using the results of proteomic analyses performed in two previous studies [30,31]. Among these proteome studies, one study used mixed samples of the seminal vesicles and sperm [30], while the other study only used sperm samples [31]. Comparing these two data sets, 66 proteins were considered candidates highly enriched in the seminal vesicles but not sperm (figure 4*a*, Table 2). Next, we browsed the *D. melanogaster* single-cell transcriptome database Fly Cell Atlas (https://flycellatlas.org/) [32] to obtain the gene expression dataset derived from the male reproductive glands. According to the Fly Cell Atlas dataset, the following four genes among the 66 candidate genes are highly enriched in the seminal vesicles as compared to other cells in the male reproductive glands (avg_logFC>2): *Lactate dehydrogenase* (*Ldh*), *Glutamate dehydrogenase* (*Gdh*), *CG10407*, and *CG10863* (figure 4*a*, Table 2). We then conducted RT-qPCR to confirm whether these genes were expressed in the seminal vesicles. The mRNA levels of all candidate genes were higher in the seminal vesicles compared to the testes and the male accessory glands (figure 4*b*-*e*).

**Figure 4.**
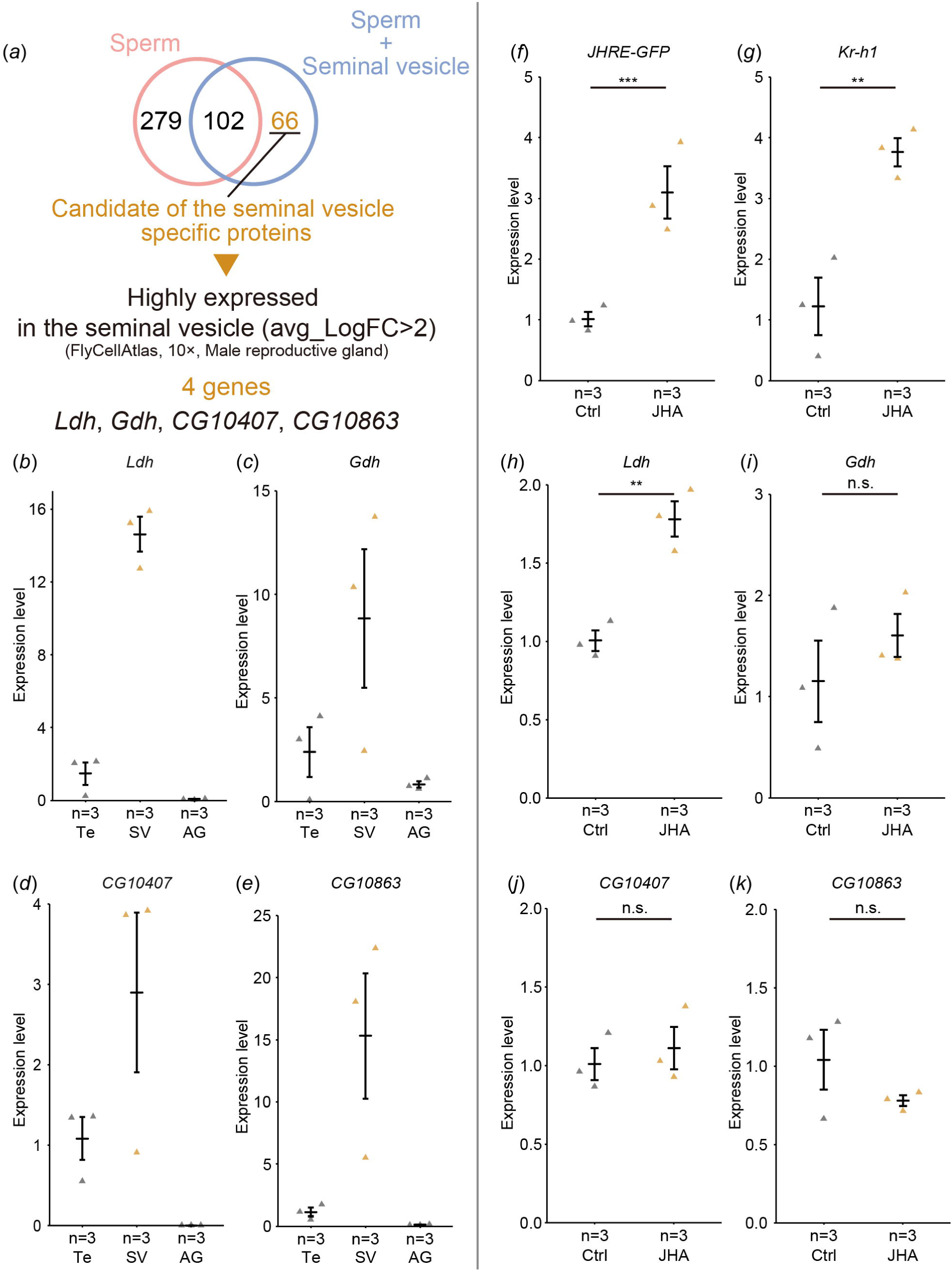
Screening of genes highly expressed in the seminal vesicle. (*a*) An overall flowchart to identify candidate genes that are highly and predominantly expressed in the seminal vesicles. See Results and Materials and Methods for details. (*b*-*e*) RT-qPCR of the candidate genes in *JHRE^WT^-GFP* males. mRNA levels were compared among the testes (Te), seminal vesicles (SV), and male accessory glands (AG). Each dot represents the levels of mRNA derived from 8 virgin males 6 days after eclosion. (*b*) *Ldh*. (*c*) *Gdh*. (*d*) *CG10407*. (*e*) *CG10863*. (*f*-*k*) RT-qPCR of the candidate genes in male reproductive tracts, including the seminal vesicles, of *JHRE^WT^-GFP* males with (JHA) or without (Ctrl) oral administration of methoprene. Each dot represents the levels of mRNA derived from 5 virgin males 7 days after eclosion. (*f*) RT-*JHRE-GFP*. (*g*) *Kr-h1*. *JHRE-GFP* and *Kr-h1* are positive control of JH responsive genes. (*h*) *Ldh*. (*i*) *Gdh*. (*j*) *CG10407*. (*k*) *CG10863*. Values in *b*-*k* are presented as mean ± SE. Statistical analysis: Student’s t-test for *f*-*k*. \*\**P* <0.01, \*\*\**P* <0.001. n.s.: not significant.

To determine whether expression of these candidate genes is regulated by JH, we did RT-qPCR to measure mRNA levels in male reproductive tracts containing the seminal vesicles dissected from *JHRE^WT^-GFP* flies with and without JHA administration. The mRNA levels of *JHRE-GFP* and the JH-responsive gene *Kr-h1*, used as positive controls, were upregulated by JHA treatment (figure 4*f*,*g*). Among the candidate genes, *Ldh* mRNA levels was upregulated by JHA treatment (figure 4*h*), while *Gdh*, *CG10407*, and *CG10863* showed no change in mRNA levels (figure 4*i*,*j*,*k*). These results suggest that JH signaling in the seminal vesicle induces the expression of *Ldh*.

To confirm whether *Ldh* is expressed in the seminal vesicle epithelial cells, we used the transgenic strain, *Ldh-optGFP*, expressing *GFP*-tagged *Ldh* under the control of *Ldh* regulatory sequences [33,34]. We found that Ldh-optGFP signal was higher in the seminal vesicles, compared with other parts of male reproductive tracts (figure 5*a*). The magnified images show that *Ldh-optGFP* is expressed in the seminal vesicle epithelial cells (figure 5*b*,*c*), suggesting that *Ldh* is highly expressed in the seminal vesicle epithelial cells. Importantly, two canonical Met/Gce binding E-box sequences (CACGTG) are found in the *Ldh* locus, one motif is located in the *Ldh-RA* promoter region and the other motif is located within the first intron (figure 5*d*). This suggests that *Ldh* is a direct target of Met and Gce. Finally, we examined whether the expression of *Ldh* was regulated by Met and Gce. We found that *Ldh* mRNA level was decreased by a double knockdown of *Met* and *gce* in the seminal vesicle epithelial cells using *Pde8-GAL4* driver (figure 5*e*). Together, these results indicate that *Ldh* is a JH-responsive gene in the seminal vesicle epithelial cells.

**Figure 5.**
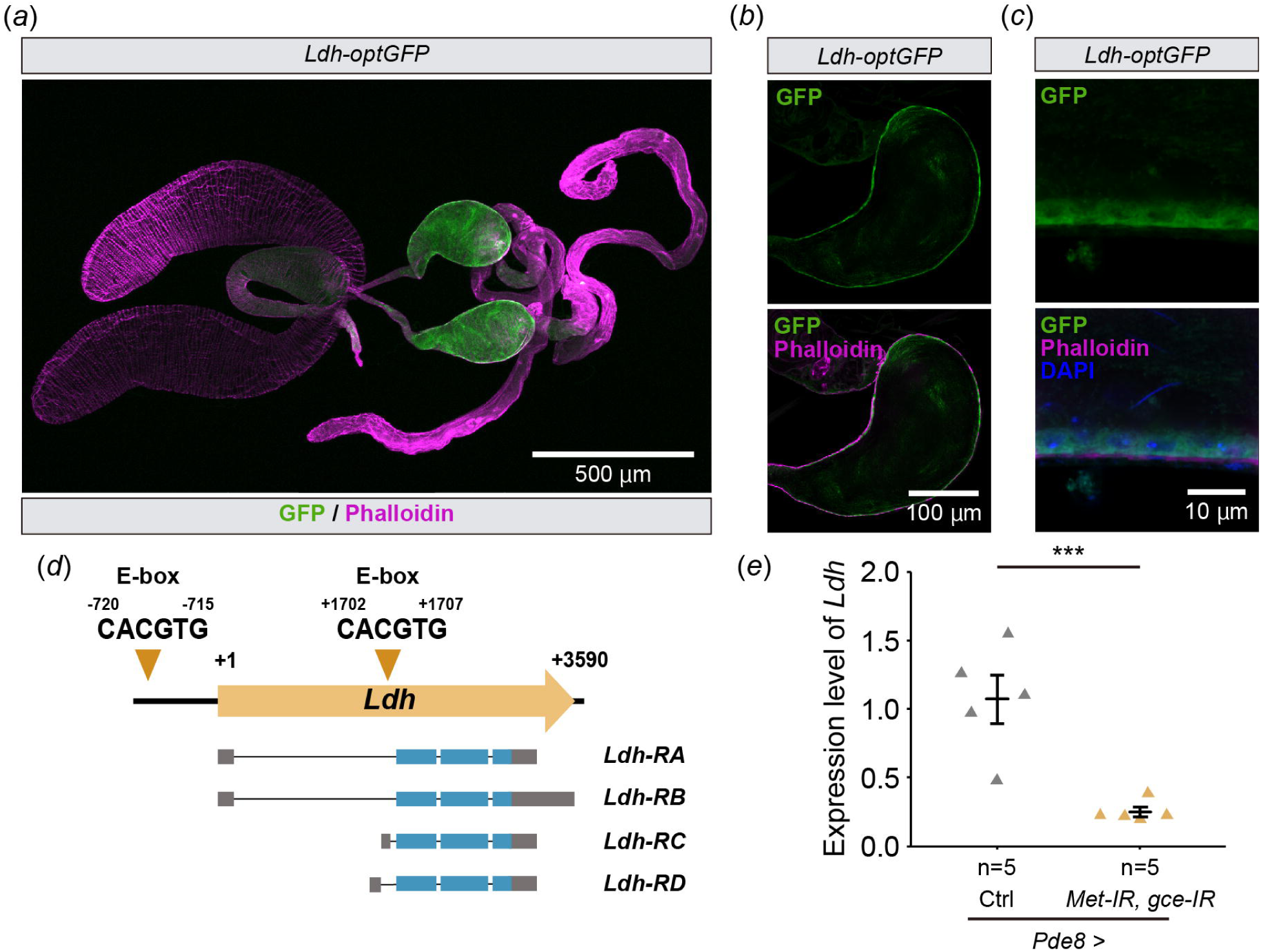
*Ldh* is expressed in the seminal vesicle epithelial cells. (*a*-*c*) Immunostaining with anti-GFP antibody (green), phalloidin (magenta), and DAPI (Blue) of *Ldh-optGFP* virgin males 4 days after eclosion. (*a*) Image of the male reproductive tract. (*b*) Cross-section image of the seminal vesicle. (*c*) Magnified view of the seminal vesicle epithelial cells. (*d*) Schematic representation of E-box in the promoter region and the first intron of *Ldh-RA*. Yellow arrow represents position of each E-box motif and numbers indicate the number of base pairs from the transcription start site (+1). Gray and blue boxes indicate untranslated and coding sequences of *Ldh*, respectively. (*e*) RT-qPCR of *Ldh* in the seminal vesicle of *Pde8-GAL4*-driven *Met* and *gce* RNAi flies. Each dot represents the levels of mRNA derived from 10 seminal vesicles of virgin males 7 days after eclosion. Values in *d* are presented as mean ± SE. Statistical analysis: Student’s t-test for *d*. ***P <0.001. n.s.: not significant.

## Discussion

In this study we identified the seminal vesicle as a JH-responsive tissue in adult male *D. melanogaster*. The seminal vesicles are present in males of many insects, including *D. melanogaster*. The seminal vesicles are known to store, nourish, and maintain sperm before they are transferred into the female reproductive tract [26]. In addition, the seminal vesicles act as secretory organs that may assist in producing seminal fluid proteins in some insects [35–40]. However, how seminal vesicles impart these functions or whether there are additional functions is not understood. In addition, neither our current study nor previous studies have been able to clarify the biological significance of the action of JH on the seminal vesicles. Considering that JH is involved in mating behavior and memory in male *D. melanogaster* [24,25,41,42], a JH-mediated modulation of the seminal vesicle’s function may also affect mating and reproduction. In fact, a previous study on the tasar silkmoth *Antheraea mylitta* has revealed that topical application of Juvenile hormone III to newly emerged adult males increases the concentration of total seminal vesicle proteins [43]. Therefore, the JH responsiveness of seminal vesicles might be evolutionarily among insects.

An important finding in this study is that the expression of *Ldh* in the seminal vesicles is upregulated by activation of JH signaling. While *Ldh* expression is known to be regulated by ecdysone signaling [44], our study is the first report that *Ldh* is also influenced by JH signaling. It is noteworthy that two canonical Met/Gce binding E-box sequences are found in the *Ldh* locus (figure 5*d*), leaving open the possibility that *Ldh* is a direct target of Met and Gce, though experimental validation of this postulate will be needed in future studies. Considering that Ldh is an essential enzyme of the anaerobic metabolic pathway [45,46], a metabolic state in the seminal vesicles might be regulated by JH. Neurobiological studies using *D. melanogaster* have shown that Ldh has an important role in supplying lactate from glial cells to neurons, known as a lactate shuttle, in response to neural activity in order to supply nutrients to neurons [47–49].

Considering the storage of many sperm in the seminal vesicles and the high expression of Ldh in the seminal vesicle epithelial cells, the lactate shuttle may exist between the sperm stored in the seminal vesicle and the seminal vesicle epithelial cells. It will be intriguing to examine whether JH signaling in the seminal vesicle changes in the quantity and/or quality of sperm.

An interesting previous study have reported that the seminal vesicle expresses multiple clock genes such as *period*, *Clock* (*Clk*), and *timeless*, all of which are necessary for generating proper circadian rhythm [50]. Considering that Met binds directly to CLK to form a heterodimer in *D. melanogaster* [51], circadian rhythm factors and JH may cooperatively regulate gene expression in the seminal vesicles.

In this study, we used both *JHRE-GFP* and *JHRR-lacZ* lines to analyze JH responsive tissues. Unexpectedly, we found that *JHRR-lacZ* and *JHRE-GFP* were differentially expressed in adult males. For example, JHRE-GFP signal was not observed in the male accessory gland, which has been reported as a JH-responsive tissue [24,52–54]. On the other hand, JHRR-LacZ signal was observed in the male accessory gland (figure 1*f*). This difference may be due to the origin of *JHRE* and *JHRR*. *JHRE* in *JHRE^WT^-GFP* strain is derived from the *early trypsin* gene of *A. aegypti* [22,23], while *JHRR* is derived from *D. melanogaster Kr-h1* [14]. Alternatively, differences in reporter activity may reflect differences in genomic context, as both *JHRE^WT^-GFP* and *JHRE^Mut^-GFP* transgenes are inserted into the *attP2* site of the third chromosome while the *JHRR-lacZ* is randomly integrated into the third chromosome. Nonetheless, activities of both reporters are restricted to a limited number of cell types of male reproductive tracts. Considering that Met and Gce are expressed in almost all cell types of male reproductive tracts [21], future studies will be needed to determine whether more comprehensive JH reporter strains are needed in *D. melanogaster* as well as other insects.

Nevertheless, we propose that the *JHRE^WT^-GFP* and *JHRE^Mut^-GFP* strains [22] are nice tools to approximate JH signaling *in vivo* in adult male seminal vesicles. For example, we found in this study that JHRE-GFP in the seminal vesicles is elevated after mating. This observation is consistent with the fact that JH titer is elevated after mating through the action of Ecdysis-triggering hormone [42]. Since *JHRE^WT^-GFP* strain has the tandem of eight JHREs [22], it may have the advantage of sensitivity for JH signaling. While direct measurements of actual JH titers are crucial [55], indirect approximation of JH titers through *JHRE^WT^-GFP* and *JHRE^Mut^-GFP* reporter activity in adult males is very easy and convenient. Use of JHRE-GFP signals in the seminal vesicles as a marker of JH signaling will facilitate future studies to increase our understanding of JH-dependent insect male physiology.

## Materials and Methods

### Drosophila melanogaster strains and maintenance

*D. melanogaster* flies were raised on a standard yeast-cornmeal-glucose fly medium (0.275 g agar, 5.0 g glucose, 4.5 g cornmeal, 2.0 g yeast extract, 150 μL propionic acid, and 175 μL 10% butyl p-hydroxybenzoate (in 70% ethanol) in 50 mL water) at 25 °C under a 12:12 hr light/dark cycle. For the JHA oral administration (figure 1*a*, 2*a*,*b*,*j*,*k*, 4*f*-*k*), virgin male flies were collected 0 to 8 hr after eclosion, aged for 4 days on standard food, and then transferred for 3 days into new tubes in the presence of food supplemented with 60 µM methoprene (Sigma-Aldrich, St Louis, MO, PESTANAL 33375, racemic mixture; 1.5 M stock was prepared in ethanol) or 0.8% ethanol (control). To analyze the effect of mating (figure 3*a*-*d*), virgin male flies were collected at eclosion, aged for 4 days on standard food and then transferred for 2 days into new tubes in the presence of *w^1118^* 4 days after eclosion virgin females. The ratio of males to females in a vial for mating was 1:2. For experiments other than JHA administration and mating, adult males were aged for 2 to 7 days on standard food.

The following transgenic strains were used: *how-GAL4* (Bloomington Drosophila stock center [BDSC] #1767), *JHAMT-GAL4* [56] (a gift from Sheng Li, South China Normal University, China), *JHRE^Mut^-eGFP* [22], *JHRE^WT^-eGFP* [22], *JHRR-LacZ* [14](a gift from Sheng Li), *KK control* (Vienna Drosophila resource center [VDRC] #60100), *Ldh-optGFP* (BDSC #94704), *Pde8-GAL4* (BDSC #65635), *UAS-GFP, mCD8::GFP* [57] (a gift from Kei Ito, University of Cologne, Germany), *UAS-gce-IR* (VDRC #101814), *UAS-jhamt-IR* (VDRC #103958), *UAS-Met-IR* (VDRC #45852), and *UAS-stinger* (BDSC #84277).

### Immunohistochemistry

The tissues were dissected in Phosphate-Buffered Saline (PBS) and fixed in 4% paraformaldehyde in PBS for 30–60 min at 25–27 °C. The fixed samples were rinsed thrice in PBS, washed for 15 min with PBS containing 0.3% Triton X-100 (PBT), and treated with a blocking solution (2% bovine serum albumin in PBT; Sigma-Aldrich #A9647) for 1 hr at 25–27 °C or overnight at 4 °C. The samples were incubated with a primary antibody in blocking solution overnight at 4 °C. The primary antibodies used were as follows: chicken anti-GFP antibody (Abcam #ab13970, 1:2,000), mouse anti-LacZ (β-galactosidase; Developmental Studies Hybridoma Bank #40-1a; 1:50). The samples were rinsed thrice with PBS and then washed for 15 min with PBT, followed by incubation with fluorophore (Alexa Fluor 488)-conjugated secondary antibodies (Thermo Fisher Scientific; 1:200) and in blocking solution for 2 hr at RT or overnight at 4 °C. Nuclear stains used in this study were 4’,6-diamidino-2-phenylindole (DAPI; final concentration 1 μg/ml Sigma-Aldrich, St. Louis, MO, USA). F-Actin was stained with Alexa Fluor 568 phalloidin (1:200; Invitrogen, #A12380). For DAPI and phalloidin staining, after the incubation with the secondary antibodies, the samples were washed and then incubated with DAPI and phalloidin for at least 20 min at RT or overnight 4 °C. After another round of washing, all the samples were mounted on glass slides using FluorSave reagent (Merck Millipore, #345789). For the quantification of JHRE-GFP signal (figure 2*a*-*f*, 3), only DAPI and phalloidin was stained after fixation. Confocal images were captured using the LSM 700 laser scanning confocal microscope (Carl Zeiss, Oberkochen, Germany). Quantification of immunostaining signal was conducted using the ImageJ software version 1.53q [58]. Fluorescence intensity of JHRE-GFP was normalized to the area of the seminal vesicle.

### *Ex vivo* male reproductive tract culture

We collected *JHRE^WT^-GFP* virgin males 4 days after eclosion. The male reproductive tracts were dissected in Schneider’s Drosophila Medium (SDM; Thermo Fisher Scientific, #21720024), and male accessory glands were removed from the male reproductive tracts using forceps. Approximately 5–6 male reproductive tracts were immediately transferred to a dish containing 3 mL of SDM supplemented with 15% fetal calf serum and 0.6% penicillin-streptomycin with/without the addition of 1 µM methoprene (Sigma-Aldrich, St Louis, MO, PESTANAL 33375, racemic mixture; 1.5 M stock was prepared in ethanol) or 0.7% ethanol (control). The cultures were incubated at 25 °C for 16 hr, and the samples were immunostained to check the JHRE-GFP signal.

### Screening of *GAL4* lines that label the seminal vesicle epithelial cells

To knock down *Met* and *gce* in the seminal vesicle, we needed a *GAL4* driver active in the seminal vesicle epithelial cells. For this purpose, we first surveyed which genes are highly and predominantly expressed in the seminal vesicles. Candidates of the seminal vesicle-specific genes were extracted from the single-cell transcriptome database, Fly Cell Atlas (https://flycellatlas.org/) [32]. In the Fly cell atlas, a transcriptomic cluster of the seminal vesicle was annotated in the 10x Genomics dataset from the whole body and the male reproductive gland samples. We extracted the gene profile of the seminal vesicle cluster derived from the whole-body sample and the male reproductive gland sample. The two profiles of gene expression datasets were filtered by *P*-value (*P*-value < 0.05) and log fold change (avg_logFC > 5). The avg_log FC indicates how specific the expression of a gene is in the certain cluster. Finally, 11 candidate genes were obtained (Table 1). Of the published *GAL4* strains under the control of each of the 11 candidates, we promptly obtained *Pde8-GAL4* and confirmed the expression pattern of *Pde8-GAL4* in the seminal vesicle as described in the main text (figure 2*g-i*).

**Table 1.**
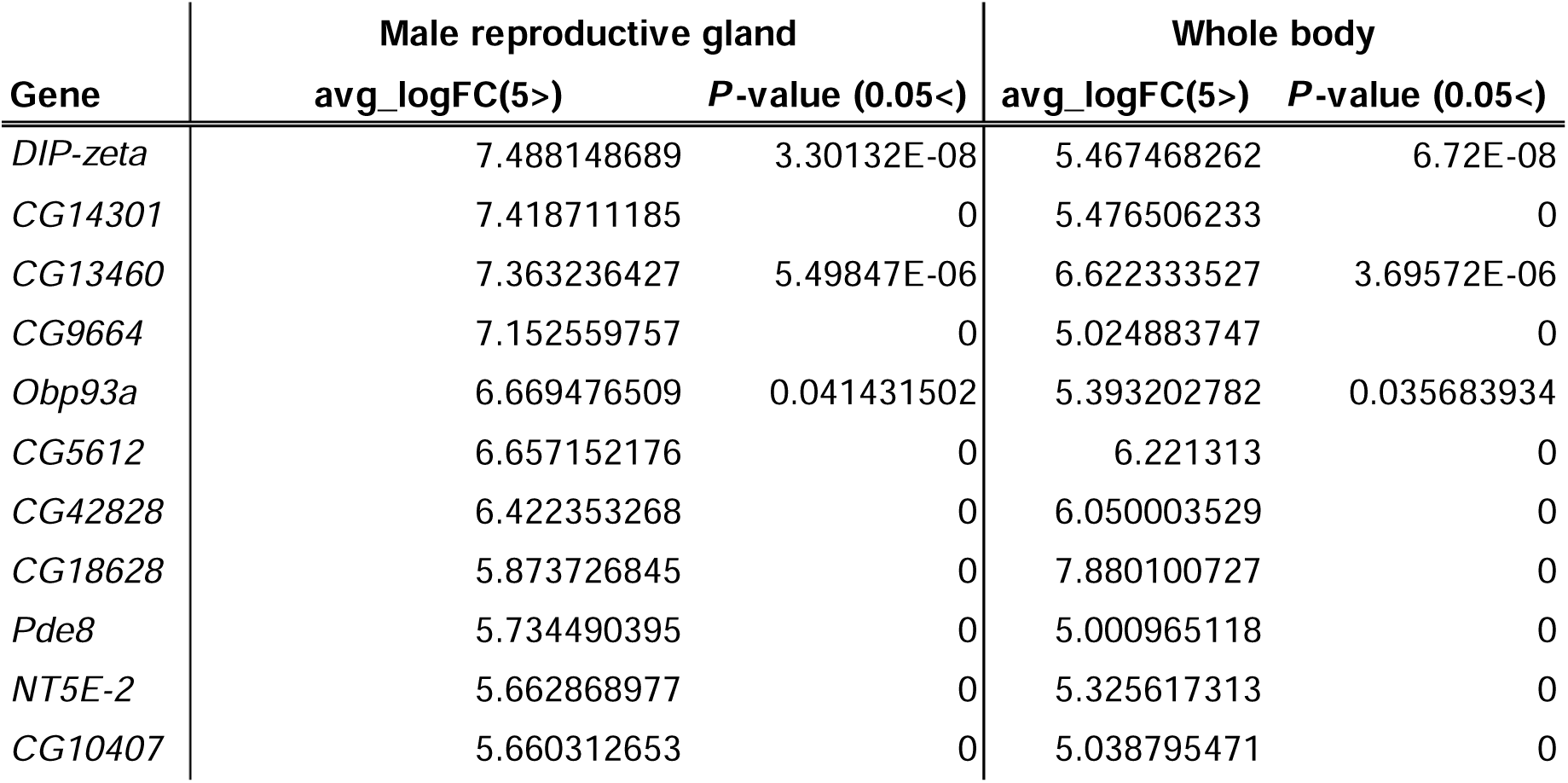
Candidate seminal vesicle-specific genes for suitable GAL4 identification. avg_LogFC: average_Log fold change

### Screening of candidate genes that are specifically and highly expressed in the seminal vesicles

Candidate proteins highly enriched in the seminal vesicle were determined by comparing the two independent proteomics datasets. One dataset [30] annotates 168 proteins as being enriched in the seminal vesicle and/or sperm stored in the seminal vesicle. Another dataset [31] annotates 381 proteins as being enriched in the sperm isolated from the seminal vesicle. We found that two datasets share 102 proteins, suggesting that these shared proteins are enriched in the sperm but not the seminal vesicle, with the remaining 66 proteins (168 minus 102) as candidate proteins enriched in the seminal vesicle (Table 2). Next, we checked whether each of the genes encoding the 66 proteins is predominantly expressed in the seminal vesicles by the single-cell transcriptome database Fly Cell Atlas (https://flycellatlas.org/)[32]. We extracted gene profiles of the seminal vesicle cluster in male reproductive gland sample. The candidate genes were filtered by *P*-value (*P*-value < 0.05) and log fold change (avg_logFC > 5). Finally, we obtained 4 candidate genes, *Ldh*, *Gdh*, *CG10407*, and *CG10863*.

**Table 2.** Candidate proteins that are specifically and highly expressed in the seminal vesicles. avg_LogFC: average_Log fold change. “Not found” indicates that the presence of mRNA in the seminal vesicle cluster could not be confirmed on the Fly Cell Atlas.

| Gene | avg_logFC<br>(Male reproductive gland) | P-value |
| --- | --- | --- |
| CG10407 | 5.660312653 | 0 |
| CG10863 | 3.085217476 | 8.75402E-11 |
| Gdh | 2.438033104 | 5.63537E-09 |
| Ldh | 2.38763833 | 2.66E-18 |
| Argk1 | 1.832550764 | 1.34915E-13 |
| regucalcin | 1.310050249 | 1.75488E-14 |
| GstD1 | 1.146826625 | 7.57E-18 |
| ldgf4 | 0.629053414 | 0.000129652 |
| Vha26 | 0.506168485 | 0.021732058 |
| awd | -0.363285989 | 0.015698655 |
| Est-6 | -0.43701005 | 1.56277E-05 |
| ldh | -0.485521317 | 0.002354908 |
| Rack1 | -0.502643883 | 0.000129842 |
| Obp44a | Not found | Not found |
| Gs1 | Not found | Not found |
| Dip-B | Not found | Not found |
| scpr-C | Not found | Not found |
| Inos | Not found | Not found |
| AdSS | Not found | Not found |
| Acsf2 | Not found | Not found |
| Bfc | Not found | Not found |
| CG11042 | Not found | Not found |
| Pglym78 | Not found | Not found |
| pyd3 | Not found | Not found |
| Pgk | Not found | Not found |
| Got2 | Not found | Not found |
| LManII | Not found | Not found |
| Fdh | Not found | Not found |
| CG8036 | Not found | Not found |
| CG3609 | Not found | Not found |
| Mfe2 | Not found | Not found |
| Chd64 | Not found | Not found |
| Irp-1B | Not found | Not found |
| Prx2540-2 | Not found | Not found |
| GstE12 | Not found | Not found |
| Gdi | Not found | Not found |
| ldgf3 | Not found | Not found |
| Cam | Not found | Not found |
| CG1648 | Not found | Not found |
| CG4520 | Not found | Not found |
| CG7264 | Not found | Not found |
| CG14282 | Not found | Not found |
| CG15125 | Not found | Not found |
| CG5177 | Not found | Not found |
| CG6287 | Not found | Not found |
| Ogdh | Not found | Not found |
| Rpi | Not found | Not found |
| CG34107 | Not found | Not found |
| ND-19 | Not found | Not found |
| ATPsynO | Not found | Not found |
| ND-B22 | Not found | Not found |
| RecQ5 | Not found | Not found |
| Fkbp12 | Not found | Not found |
| Swim | Not found | Not found |
| Mp20 | Not found | Not found |
| Tm2 | Not found | Not found |
| Tm1 | Not found | Not found |
| Prm | Not found | Not found |
| Mf | Not found | Not found |
| Mhc | Not found | Not found |
| Tsf1 | Not found | Not found |
| Zasp66 | Not found | Not found |
| porin | Not found | Not found |
| PPO1 | Not found | Not found |
| Sod2 | Not found | Not found |
| CG11815 | Not found | Not found |

### Reverse transcription-quantitative PCR (RT-qPCR)

RNA from tissues was extracted using RNAiso Plus (Takara Bio) and reverse-transcribed using ReverTra Ace qPCR RT Master Mix with gDNA Remover (TOYOBO). Synthesized cDNA samples were used as templates for quantitative PCR using THUNDERBIRD SYBR qPCR Mix (TOYOBO) on a Thermal Cycler Dice Real Time System (Takara Bio). The amount of target RNA was normalized to the endogenous control *ribosomal protein 49* gene (*rp49*) and the relative fold change was calculated. The expression levels of each gene were compared using the ΔΔCt method [59]. The following primers were used for this analysis: rp49 F (5’-CGGATCGATATGCTAAGCTGT-3’), rp49 R (5’-GCGCTTGTTCGATCCGTA-3’), GFP F (5’-GAACCGCATCGAGCTGAA-3’), GFP R (5’-TGCTTGTCGGCCATGATATAG-3’), CG10407 F (5’-ACTGGACAACAGCCAAACCTC-3’), CG10407 R (5’-GTGTCTAGGTCGGGTGCATTG-3’), Ldh F (5’-CGTTTGGTCTGGAGTGAACA-3’), Ldh R (5’-GCAGCTCGTTCCACTTCTCT-3’), Gdh F (5’-GGAGGACTACAAGAACGAGCA-3’), Gdh R (5’-CAGCCACTCGAAGAAGGAGA-3’), CG10863 F (5’-CATCGGACTGGGCACCTATAC-3’), CG10863 R (5’-TTCTCGTAGAAATAGGCGGTGTC-3’), Kr-h1 F (5’-TCACACATCAAGAAGCCAACT-3’), Kr-h1 R (5’-GCTGGTTGGCGGAATAGTAA-3’).

### Statistical analysis

All experiments were performed independently at least twice. The sample sizes were chosen based on the number of independent experiments required for statistical significance and technical feasibility. The experiments were not randomized, and the investigators were not blinded. All statistical analyses were performed using the “R” software version 4.0.3. Details of the statistical analyses are described in figure legends.

## Acknowledgments

We thank Sheng Li, Kei Ito, Naoki Yamanaka, Bloomington Stock Center, Vienna Drosophila Resource Center for fly strains, Developmental Studies Hybridoma Bank for antibodies, and Jason Tennessen, Daiki Fujinaga, Ryo Hoshino, Eisuke Imura, Yuto Yoshinari, and Yoshiki Hayashi for helpful discussions.

## Author contributions

YK: Conceptualization, Validation, Formal analysis, Investigation, Data Curation, Writing – Original Draft, Visualization, Funding acquisition

YM: Validation, Investigation, Writing – Review & Editing, Funding acquisition

NO: Methodology, Resources, Writing – Review & Editing

LB: Methodology, Resources, Writing – Review & Editing

RN: Conceptualization, Resources, Writing – Original Draft, Visualization, Supervision, Project administration

